# Dissociating External and Internal Attentional Selection

**DOI:** 10.1101/2024.08.27.609883

**Authors:** Kabir Arora, Surya Gayet, J. Leon Kenemans, Stefan Van der Stigchel, Samson Chota

**Affiliations:** Helmholtz Institute, Utrecht University, 3584 CS, Utrecht, The Netherlands

## Abstract

Visual Working Memory (VWM) stores visual information for upcoming actions. Just as attention can shift externally towards relevant objects in the visual environment, attention can shift internally towards (i.e., prioritize) VWM content that is relevant for upcoming tasks. Internal and external attentional selection share a number of key neural and functional characteristics, which include their spatial organization: recent work has shown that spatial attention is directed towards the previous location of a prioritized memory item, similar to how a perceived stimulus is prioritized. Attending stimuli that are physically present is useful, as it enhances processing of the relevant visual input. When prioritizing items in memory, however, attending the prior stimulus location cannot serve this purpose, as there is no visual input to enhance. Here, we address this apparent contradiction which highlights the gaps in our understanding of the mechanisms underlying external and internal visual attention. In two EEG experiments, we compare location-specific sensory enhancement during the attentional selection of external (perceived) as compared to internal (memorized) stimuli. During both internal and external selection we observed a lateralization of alpha oscillations and gaze position bias toward the previous locations of prioritized items, confirming earlier findings that suggested an inherent spatial organization within VWM. Critically, using Rapid Invisible Frequency Tagging (RIFT), we show that sensory enhancement at the attended location is only observed during external attentional selection of (perceived) stimuli. No such location-specific sensory enhancement was observed during attentional selection of items in VWM. Furthermore, we found no clear relationship across trials between alpha lateralization and sensory enhancement (measured through RIFT) during external attention, suggesting that these two metrics indeed reflect distinct cognitive mechanisms. In sum, using a novel combination of EEG and RIFT, we demonstrate a fundamental distinction between the neural mechanisms underlying the selection of perceived and memorized objects. Both types of selection operate within a spatial reference frame, but only external selection modulates early sensory processing. Our findings suggest that the visual system is not vestigially recruiting existing mechanisms of external attention for prioritization in VWM, but is instead using space as an organizational principle to store and select items in VWM.

## Introduction

Visual Working Memory (VWM) enables the temporary maintenance of visual stimuli to facilitate perception and upcoming actions. When multiple items are held within VWM, one item might be more relevant than others for an upcoming task. Just as you can shift attention externally to select relevant objects in the visual environment, you can shift attention internally to relevant content in VWM. For example, as you approach the confectionery aisle of the grocery store, you may prioritize the mental representation of your favorite brand of cookies, as opposed to that of milk or bread. Although external attention (prioritizing currently available visual input) and internal attention (prioritizing VWM information that is no longer available externally) are two seemingly different processes, it has been shown that internal attention shares some neural and functional markers of external attention (Griffin & Nobre, 2003; Kiyonaga & Egner, 2013; Nobre et al., 2004). Despite such markers of similarity, it is currently unclear to which extent internal and external attentional selection recruit the same neural mechanisms; does our hyperefficient neural system reflect this similarity by consolidating the mechanisms the two use into one shared pipeline?

The external world is inherently spatially organized; different objects occupy different locations in our environment. During external selection, we may thus direct our spatial attention to locations with relevant stimuli, thus boosting the processing of visual input at the corresponding locations. Somewhat surprisingly, evidence suggests a similar reliance on location during selection in VWM, even when there is no relevant information at these locations anymore. For example, when asked for the color of your lost phone, you may automatically direct attention to the pocket in which you usually keep it while retrieving its color from memory.

Evidence for a spatial organization in VWM mainly stems from studies using retro-cue paradigms. A retro-cue is presented after a set of items has already been memorized, and informs the participant about the relevance of an item within this set for an upcoming task. Recall performance is better for retrocued items, suggesting that the cue elevates the item to a prioritized state within VWM (Souza & Oberauer, 2016). It has been shown that retroactively cueing an item in VWM produces a behavioral bias towards the location where this item was previously memorized, even if memorizing item locations was not necessary to complete the task. Reaction times to perceptual targets that are displayed following a retro-cue are faster when they match the location where the retro-cued item was memorized (Awh et al., 1998; Theeuwes et al., 2011). In similar tasks, retro-cues induce microsaccadic eye movements towards the previously memorized item location (Van Ede et al., 2019), as well as desynchronization of alpha power contralateral to the encoding location of the item (Liu et al., 2022; Poch et al., 2017; Sauseng et al., 2009). Alpha lateralization is an established neural signature of orienting visual attention (Ikkai et al., 2016; Kelly et al., 2006; Thut et al., 2006; Worden et al., 2000). Taken together, this work shows that even if the former encoding locations of memory items are no longer important (they contain no task-relevant sensory information), maintenance and selection of items in VWM still seem to operate within a spatial layout that mirrors the layout of our external world.

Despite clear evidence for a spatial organization in VWM, there is one key distinction to be made between such a spatial organization in external and internal attention. Attentional shifts towards (external) perceived stimuli serve the encoding of task-relevant sensory input at that location. In contrast, when a retro-cue instructs participants to favor one stimulus over the other, the former location of the memorized stimulus no longer contains any relevant information. Thus, although the literature reviewed above showed an attentional bias toward the former location of retro-cued stimuli, VWM cannot benefit from this spatial organization in the same way that external attention does. During external attention, excitability of the earliest sensory cortices are boosted at these locations in order to facilitate the encoding of sensory information (Posner & Gilbert, 1999; Zhigalov et al., 2019). Although internal attention also produces (behavioral and neural) biases toward the attended location (Liu et al., 2022; Poch et al., 2017; Sauseng et al., 2009; Theeuwes et al., 2011; Van Ede et al., 2019), this does not automatically imply that these biases are similarly accompanied, or caused, by an enhancement of such early sensory processing. There is potential support both for the presence and absence of this early sensory enhancement during internal attention. Features of internally prioritized items can be recovered from sensory areas (Munneke et al., 2012), suggesting that they are maintained in sensory regions. As these early visual processing regions are inherently retinotopically organized, one might expect that the prioritization of visual memories also leads to a facilitation of the early sensory response to external input at former encoding locations. From a functional point of view, however, such a boost would not serve its traditional advantage of encoding external sensory information, since there is none at these locations. Here, we resolve this intriguing contradiction and test whether internal attention leads to location-specific sensory enhancement just like selecting objects externally (Figure 1.1).

**Fig 1.1:**
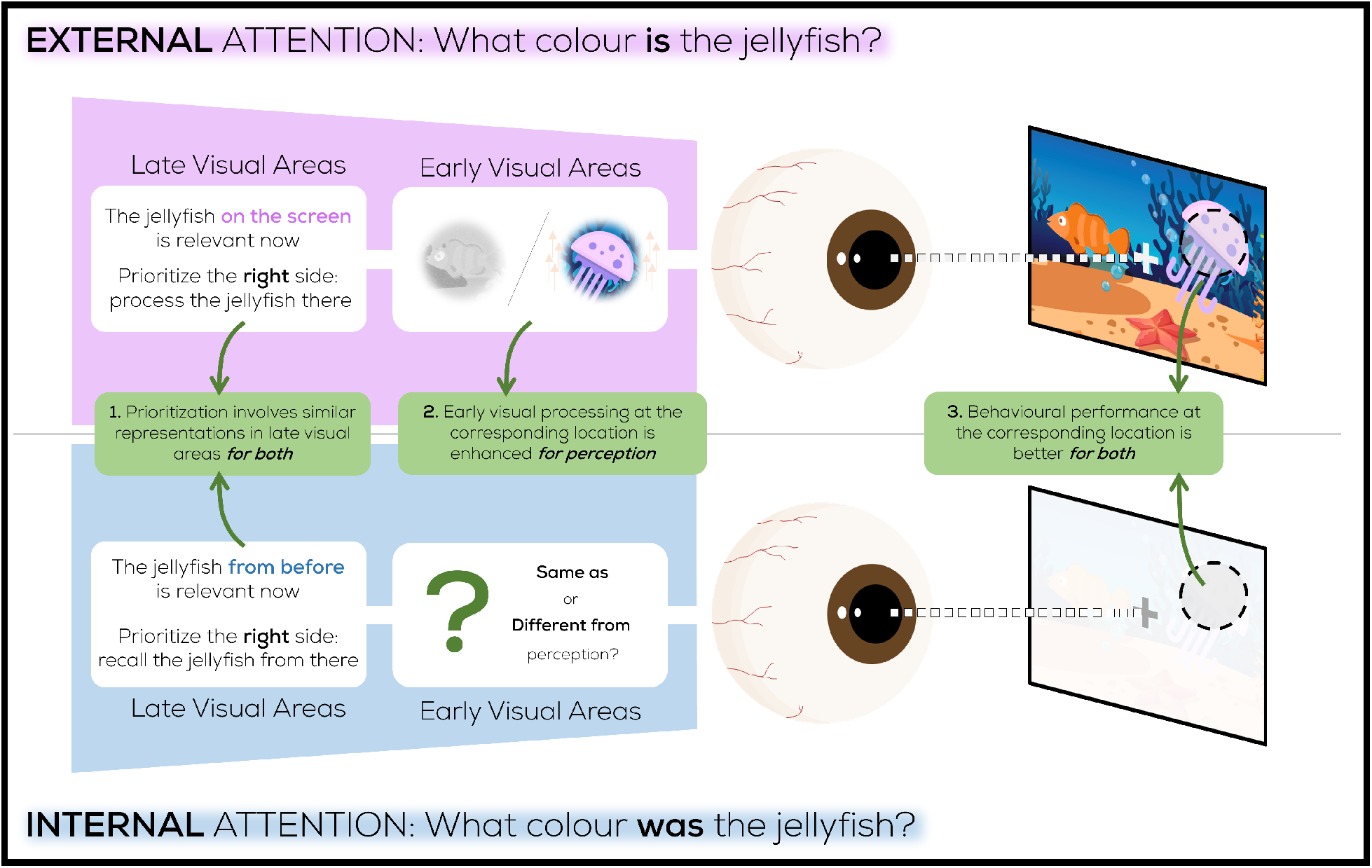
Similarities between the representation of location during internal (blue region) and external (purple region) prioritization exist in high-level visual areas (left), and behavioral performance at the corresponding location improves for both (right). It is currently unknown whether early visual processing behaves similarly across both. This may differ because of the different functions involved encode novel stimuli vs. highlight memorized information. A distinction at the early processing level could indicate whether internal attention is vestigially using spatial mechanisms that evolved for processing the external world, or that this spatial layout within VWM serves a functional purpose.

We want to measure whether sensory processing is boosted at specific locations. However, presenting a perceptible stimulus at the corresponding location would interfere with this measurement. This is because in that case, altered responses to a stimulus would not imply that the probed location would have also been processed differently in the absence of the stimulus. The perceptible stimulus may change the attentional priority of the location, or stimulus processing may propagate to downstream regions affected by attention, even if sensory processing was not boosted at its location. Though this may appear to be an obstacle since visual processing is conventionally studied with perceptible stimuli, recent methodological advances have made it possible to drive early visual cortex without usual contributions from downstream visual processing regions associated with conscious perception. Rapid periodic stimulation induces a rhythmic response that predominantly emerges from early sensory areas (Duecker et al., 2021; Ferrante et al., 2023). A recent technique known as Rapid Invisible Frequency Tagging (RIFT) (Zhigalov et al., 2019) leverages this form of stimulation to offer a measure of early visual processing at specific locations. In doing so, it forms an invisible tracker of spatial attention. Here, we use RIFT to measure the extent to which early visual processing at locations previously or currently occupied by stimuli is modulated during both internal and external shifts of attention. We show that despite the present of clear correlates of attention (a lateralization of alpha oscillations and gaze position), early visual processing is not enhanced at the previous locations of memorized items (i.e., during internal attention), even though this increase is seen at cued item locations during external attention. We discuss the implications of our results for the mechanism underlying attentional prioritization in VWM, and how they relate to the organizational principles of VWM.

## Methods

### Participants

We recruited a total of 48 healthy participants (38 female, 22.7 ± 2.6 years old; mean ± std) with normal or corrected-to-normal vision for two separate experiments. None of the participants reported a history of epilepsy or psychiatric diagnosis. Participants were compensated either with €20 or an equivalent amount of participation credits as per Utrecht University’s internal participation framework. The study was carried out in accordance with the protocol approved by the Faculty of Social and behavioural Sciences Ethics Committee of Utrecht University.

### Protocol

Participants underwent a 2-hour experimental session at the Division of Experimental Psychology, Utrecht University. Participants received procedural information prior to the session, and provided informed consent, date of birth, biological sex, and dominant hand information at the beginning of the session. After completion of the EEG setup, participants were seated 76cm from the screen with a chinrest. After eye-tracker calibration, the experiment was explained using a visual guide and verbal script. Participants were also informed that they may potentially see visual glitches or flickers on the screen (due to the high-speed projection and tagging, *see Tagging Manipulation*), and that they would be asked after the experiment whether they did or not. Following these instructions and 10 practice trials, the ∼ 1 hour task was completed. Participants filled out a questionnaire on whether they noticed any visual artifacts on screen (and if so, at what stage of the task, and to what degree they felt this interfered with their task on a scale of 1-5), which confirmed that tagging was difficult to perceive (*see Tagging Manipulation*). Compensation was awarded when applicable, and the session was ended.

### Experimental Design and Procedure

Two oriented and uniquely colored (blue or red) gratings were presented on both sides of fixation, and participants performed a delayed orientation change detection task on one of them as informed by a color cue.

There were two different experiments; one in which the cue was presented after the gratings (retro-cue task; Figure 2.1; top), and one in which the cue was presented prior to the gratings (pre-cue task; Figure 2.1; bottom).

**Fig 2.1:**
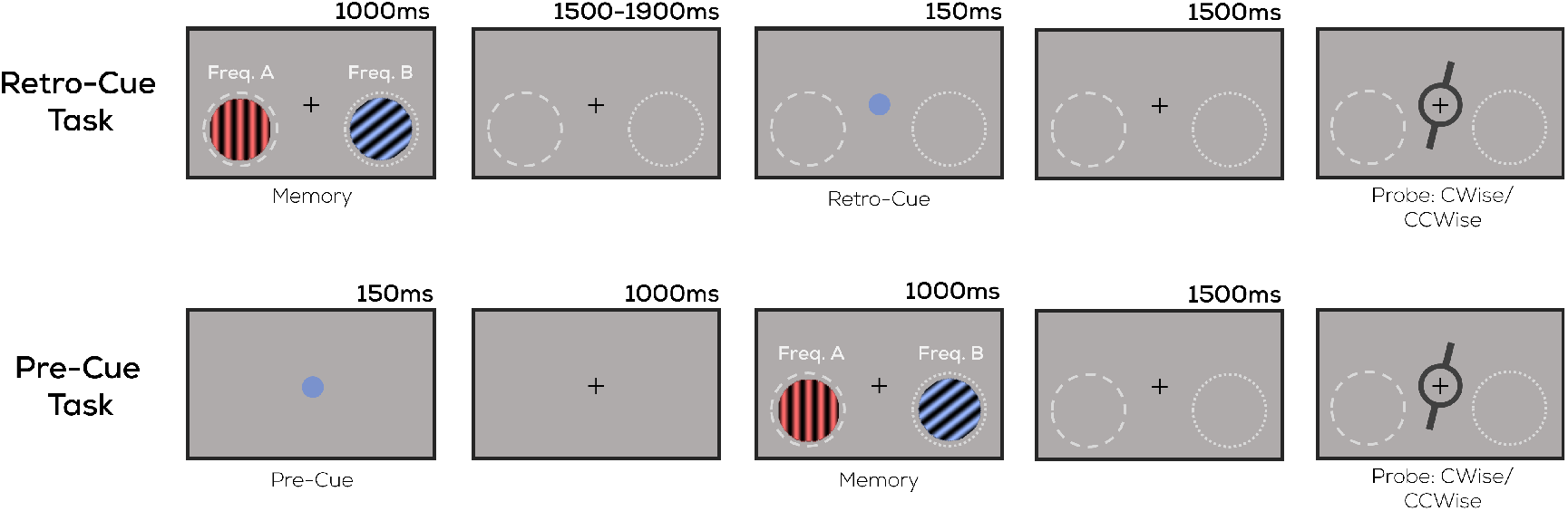
Task Design. Two uniquely colored and oriented stimuli were presented on either side of fixation. This was followed by (**retro-cue task**) or preceded by (**pre-cue task**) a red or blue circle (the cue), indicating which of the two oriented stimuli (i.e., the red or blue item) would be probed. After a delay, participants indicated whether a memory probe was tilted clockwise or counterclockwise relative to the cued item. The highlighted locations flickered at 60 or 64Hz (*See Tagging Manipulation*) throughout the trial (Note: not to scale; memory probe did not overlap with flickering regions in actual display. No actual outlines around flickering regions were displayed).

## Encoding and Cue

### Retro-Cue Task

Trials began with a centrally presented fixation cross (uniformly random duration between 1-1.25s), after which both memory items (gratings) were presented (1s). This was followed by a variable delay (uniformly random, 1.5-1.9s) after which a central retro-cue was presented (0.15s) in the form of either a blue or red circle, informing participants which item would be probed on that trial.

### Pre-Cue Task

Trials began with a centrally presented fixation cross (uniformly random duration between 1-1.25s), after which a central cue was presented (0.15s) in the form of either a blue or red circle, informing participants which upcoming item would be probed on that trial. This was followed by a fixed delay (1s), at the end of which both memory items were presented (1s).

Beyond this crucial difference in the order of cue-grating presentations, the retro-cue and pre-cue tasks were identical.

### Report

After a fixed cue-probe or grating-probe interval (1.5s), the memory probe was displayed centrally until response. Participants had to specify with a keyboard button press whether the orientation of the memory probe was more clockwise (Q key) or anti-clockwise (P key) compared to that of the cued memory item. Participants were able to respond no sooner than 0.5s after the memory probe onset (indicated via a change in the color of the fixation cross from black to green). Lastly, feedback in the form of a green check mark or red cross was given for correct or incorrect trials respectively. If no response was received within 3s, or a different button was pressed, the trial ended (median = 0.2% of trials, mean = 0.36% of trials).

Each participant completed 480 trials (excluding 10 practice trials, not included in analysis) divided into 15 blocks of 32 trials each. The retro-cue/pre-cue that was blue or red instructed participants to compare the probe orientation to either the orientation of the blue or the red memory item respectively. Orientations of the memory items (always distinct), as well as whether the blue/red stimuli would be presented to the left or right, were equated in prevalence separately and presented in random order. The location of the tagging frequencies (60L/64R vs. 64L/60R) and the direction (left/right) of the cued item were counterbalanced and presented in a random sequence.

### Stimuli

The screen background was maintained at gray (grayscale: 127.5) throughout the experiment. A black fixation cross (0.4 degrees of visual angle; dva) was present in the center of the screen throughout each trial. Memory stimuli consisted of circular square wave gratings (*r* = 3 dva, spatial freq. = 2 cpdva) either blue or red in color. These were presented slightly below horizontal (eccentricity = 6 dva horizontal, -2 dva vertical) in order to facilitate the tagging response (Minarik et al., 2023). To remove visible edges at the boundary of the tagged region, a radially symmetric transparency mask was applied to the memory stimuli as per Equation (1):

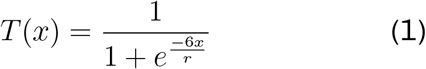

In Equation (1) *x* is the distance of a point on the circular patch from center (0) to the circumference (at radius *r*), and *T* (*x*) is the resulting transparency at this point ranging from 0.5 (semi-transparent) to 1 (fully opaque). Both memory stimuli were presented with distinct orientations ranging from 0 to 165° in intervals of 15° (thus covering the full range of possible orientations in discrete steps). The cue in both tasks (*r* = 0.23 dva) was displayed centrally in one of the two memory stimuli colors. The memory probe consisted of a black, circular annulus (*r* = 1.2 dva, thickness = 0.13 dva) with spokes (0.6 dva extensions) protruding outwards from diametrically opposite ends to indicate an orientation. These dimensions ensured that the tips of the memory probe never overlap (minimum separation = 1.5 dva) with the flickering regions on screen (*See Tagging Manipulation*). The orientation of the memory probe could vary between 3° and 50° clockwise or anticlockwise relative to the cued memory stimulus, following a staircase procedure (PsychtoolBox QUEST algorithm: (Farell & Pelli, 1999)) that maintained accuracy on the memory task at 75% (*β* = 3.5, Δ = 0.01, *γ* = 0.5).

### Tagging Manipulation

We used Rapid Invisible Frequency Tagging (RIFT) to evoke an oscillatory response from specific locations on the screen (Seijdel et al., 2023; Zhigalov et al., 2019). This involved sinusoidally varying the screen luminance at certain locations at specific frequencies. The areas corresponding to the two memory items were tagged from grating onset until the end of the trial. Two frequencies (60Hz and 64Hz) were randomly assigned to either the left or right area, resulting in two possible configurations: 60 left, 64 right; or 64 left, 60 right. When memory stimuli were presented, the two circular gratings were tagged with the corresponding frequency. For the remainder of the trial, the background at the two stimuli locations was tagged (from white to black in order to look invisible against the gray background). The same transparency map was applied to the flickering regions as the memory stimuli (*See Stimuli*). The tagging sinusoids were phase locked to cue onset in both experiments. Prior to data collection, the displayed tagging frequency was verified using a BioSemi PhotoCell luminance sensor (BioSemi B.V., Amsterdam, The Netherlands). Temporal precision of the displayed stimuli was continually recorded during data collection using PsychToolBox’s Screen(‘Flip’) command. Any trial with a frame displayed >4ms off-time was excluded from analysis (median = 0% of trials, mean = 0.16% trials). Given recent research that has established the subjective undetectability of RIFT (Spaak et al., 2024), as well as the fact that out of 48 participants in the current study only 4 reported seeing any abnormality on the screen, of which only 1 reported (low-moderate) distraction from the task, we concluded that our tagging was sufficiently difficult to perceive.

### Display Apparatus

Stimuli were projected using a ProPixx projector (VPixx Technologies Inc., QC Canada; resolution = 960×540px; refresh rate = 480Hz) in a rear-projection format (projected screen size = 48×27.2cm). Experimental code was written in MATLAB (MATLAB, 2022), using PsychToolBox3 (Brainard & Vision, 1997) for task display.

### EEG Recording and Pre-processing

EEG data was recorded using a 64-channel ActiveTwo BioSemi system (BioSemi B.V., Amsterdam, The Netherlands) at 2048Hz. Two additional electrodes were placed above and on the outer canthus of the left eye respectively. Immediately prior to the experiment, adequate signal quality from all channels was ensured using BioSemi ActiView software. All data analysis was conducted in MATLAB using the Fieldtrip toolbox (Oostenveld et al., 2011). The EEG data was first rereferenced to the average of all channels (excluding poor channels determined by visual inspection, median = 11 [frontal] channels, mean = 10.7 channels). Data was high-pass filtered (0.01Hz), then line noise and its harmonics were removed using a DFT filter (50, 100, 150Hz). Data was segmented into trials ranging from 3.4s before to 2s after retro-cue onset (retro-cue task), or 2s before to 2.5s after stimuli onset (pre-cue task). An ICA was performed to remove oculomotor artifacts, and trials with other motor artifacts were removed from further EEG analysis as per visual inspection (median = 14.3%, mean = 14.9%). Baseline correction was performed by averaging (and then subtracting from the signal) a window 0.8s to 0.3s before memory stimuli onset in the retro-cue task, and 0.8s to 0.1s before cue onset in the pre-cue task. Two participants (from the retro-cue task) with excessive noise (>50% trials labeled as artifacts) were excluded from further EEG and eye-tracking analysis.

### RIFT Response and Coherence

In order to determine the strength of the EEG response to RIFT frequencies, magnitude-squared coherence was used, which is a dimensionless quantity (ranging from 0-1) that measures how consistently similar two signals are in both their power and phase. This results in higher values when two signals i) oscillate at the same frequency, and ii) maintain the same phase difference across trials (i.e., oscillatory responses across successive trials are consistently phase-locked). Coherence was computed between a reference wave (pure sinusoids with the corresponding frequency of 60Hz or 64Hz, sampled at 2048Hz) and condition-specific sets of trials per channel and participant. Segmented trials were first bandpass filtered (*±* 1.9Hz) at the frequencies of interest (60Hz & 64Hz) using a two-pass Butterworth filter (4th order, hamming taper). The filtered time-series data was Hilbert transformed. This provided a time-varying instantaneous magnitude (*M*(*t*)) and phase (*ϕ*(*t*)). The set of all instantaneous magnitudes of the filtered responses 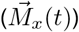 and the reference sinusoid 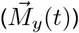 across all *n* trials, as well as the differences between their instantaneous phases across all *n* trials 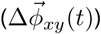 were used to compute time-varying coherence.

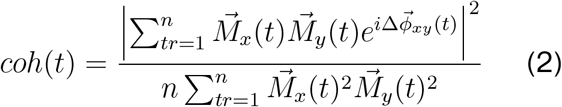

In order to compute coherence spectrograms, Equation (2) was computed for frequencies ranging from 56.8Hz to 67.2Hz in 0.8Hz intervals. We identified the top 6 channels per individual (Pan et al., 2021; Husta et al., 2024, preprint) with the strongest coherence at 60/64Hz across all trials, and any further comparisons across experimental conditions were made using traces averaged across these channels.

### Alpha Lateralization

Alpha lateralization is an established neural signature of orienting visual attention. We thus investigated whether alpha (∼ 10Hz) oscillations decreased contralateral to the hemifield where the cued item was encoded (retro-cue task) or displayed (pre-cue task). Time-Frequency representations were computed using the ft_freqanalysis function in the Fieldtrip toolbox (Oostenveld et al., 2011). First, spectral analysis was performed on individual trials in the 8-13.5 Hz range (in increments of 0.2Hz). This was done for every channel using 3-cycle Morlet wavelets and baselined (dB; 10*log10(signal/baseline)) with respect to a window 0.6s to 0.3s before memory stimuli onset (retrocue task) or before pre-cue onset (pre-cue task). The 8-13.5Hz frequency range was then averaged to reflect alpha power, and used to assess the differences in alpha band activity between conditions for each channel. Lateralization of alpha power was visualized and quantified by subtracting the alpha power of left-cued trials from right-cued ones.

### Eye-tracking Recording and Analysis

Gaze was tracked using an Eyelink SR (SR Research, Ontario, Canada) eye-tracker. Both eyes were tracked at 500Hz. Immediately prior to the experiment, a 9- point calibration was performed. This calibration was repeated after every 3rd experimental block.

The data was segmented into trials of -1.5s before to 1.5s after retro-cue onset (retro-cue task) or stimuli onset (pre-cue task). Blink correction was carried out using custom code adapted from existing work (Hershman et al., 2018). Trials were baseline-corrected with respect to the average position in an 800ms window prior to retro-cue or stimuli onset (in the retro-cue and pre-cue tasks respectively). Trials where fixation was not maintained (defined as gaze being further than 2 dva from fixation for more than 50ms of the trial duration) or trials where the baseline was over 2 standard deviations away from the mean baseline position were removed from further eye-tracking analysis (median = 17.9% of trials, mean = 18.9% of trials). The EEG results presented here do not exclude all these trials, however, we separately ran the EEG analysis without these trials which resulted in the same results qualitatively. Three participants (retro-cue task) were excluded from gaze position bias analysis for failing to maintain fixation in more than half of the trials. A 10ms uniform smoothing filter was applied to the individual position data.

### Statistical Analysis

Differences in time-varying measures (coherence and gaze position) across conditions were compared over the duration of a trial using a non-parametric cluster-based permutation test. Coherence traces were first averaged across the top 6 channels per individual with the strongest RIFT coherence at 60/64Hz. This produced a single coherence trace over time per participant for each condition and each frequency. Similarly, both eyes were averaged to produce a single horizontal bias trace over time per participant per condition. Then, a permutation test (Maris & Oostenveld, 2007) was used to inspect differences across attentional conditions. This consisted of four steps. 1) Two traces being compared were subtracted to produce a difference trace. 2) A one-sample t-test was run for each individual time point to detect points on the difference trace significantly different from 0 (*p* < 0.05). Clusters of consecutively significant timepoints longer than 10ms were identified and their sum of *t*-values was computed within each cluster to produce a cluster-level t-mass. 3) Then, we randomly flipped the sign of the different values at half of the timepoints. We conducted 10,000 repetitions of this process to generate a distribution of expected t-mass values given randomized labels. 4) Finally, we checked whether the t-masses of any initially observed clusters were higher than 95% of this distribution. These clusters were accepted as significant. With coherence, we compared the RIFT response evoked from cued vs. uncued item locations first separately for both frequencies, and later averaged. For gaze, we compared horizontal gaze position for cued left vs. cued right trials.

To compare the degree of attentional modulation in the RIFT response and the lateralization of alpha oscillations between the retro-cue and pre-cue tasks, we used a permutation test of mean differences (Ho et al., 2019). Given two sets of data points (here, one value per participant from the retro-cue and the precue tasks), this test uses bootstrap resampling to draw a distribution of the difference between the means of the two sets. On each of its 5000 repetitions, it samples points (with replacement) from both sets, and computes the difference in the means of these two sample sets. The two sets of data points are accepted as significantly different if the 95% confidence interval of this distribution excludes 0.

### Linear Mixed-effect Modeling

We lastly used a Linear Mixed-effects Model (LMM) to assess whether the attentional modulation found in the RIFT response could be explained by the other two effects observed, namely alpha lateralization and the gaze bias towards the cued item. LMM analysis was conducted in MATLAB (MATLAB, 2022), using the <monospace>fitglme</monospae> function. The corresponding formula used was RIFT Response = 1 + Cued/Uncued + Frequency (60/64) + Gaze Position Bias + Alpha Lateralization + (1 + <all predictors> | Participant). For each recorded metric, a trial-wise, time-averaged value was obtained by averaging the significant interval at the group level as determined by the cluster-based permutation tests. Trial-wise alpha lateralizations were computed using channels significant at the group level, by subtracting the ipsilateral from the contralateral alpha power on each trial. For the RIFT response in the retro-cue task, where no attentional modulation was found, we used the significant interval from the RIFT response in the pre-cue task. A second LMM assessed whether alpha lateralization could be explained by the other effects. The corresponding formula used was Alpha Lateralization = 1 + RIFT Response + Gaze Position Bias + (1 + <all predictors>|Participant). In both cases, to obtain trialwise measures of the RIFT response, we used the trialwise magnitude resulting from the Hilbert transform of the EEG signal (*See* ‘*RIFT Response: Coherence*’).

## Results

We investigated whether internal attention (prioritizing relevant content in VWM) boosts early visual processing, similar to external attention (orienting towards relevant stimuli in the environment) by running two corresponding experiments.

Behavioral results from both tasks showed that participants performed the task successfully. Performance was above chance on the pre-cue task (mean = 83.4%, median = 85.2%) as well as the retro-cue task (mean = 76.7%, median = 78.3%). All further analysis shown here was conducted using all trials, however, we separately analyzed the data using only correct trials and observed the same results qualitatively.

We first show that our tagging manipulation successfully evoked an oscillatory response in the EEG signal. Coherence (from individuals’ top 6 channels, *see Statistical Analysis*) showed a clear peak at the tagged frequencies (Figure 3.1A) and was highest around parietal/occipital electrodes (Figure 3.1B) in both tasks as expected from previous RIFT studies (Duecker et al., 2021; Ferrante et al., 2023), showing that we can successfully pick up two simultaneous RIFT-tagged responses using EEG.

**Fig 3.1:**
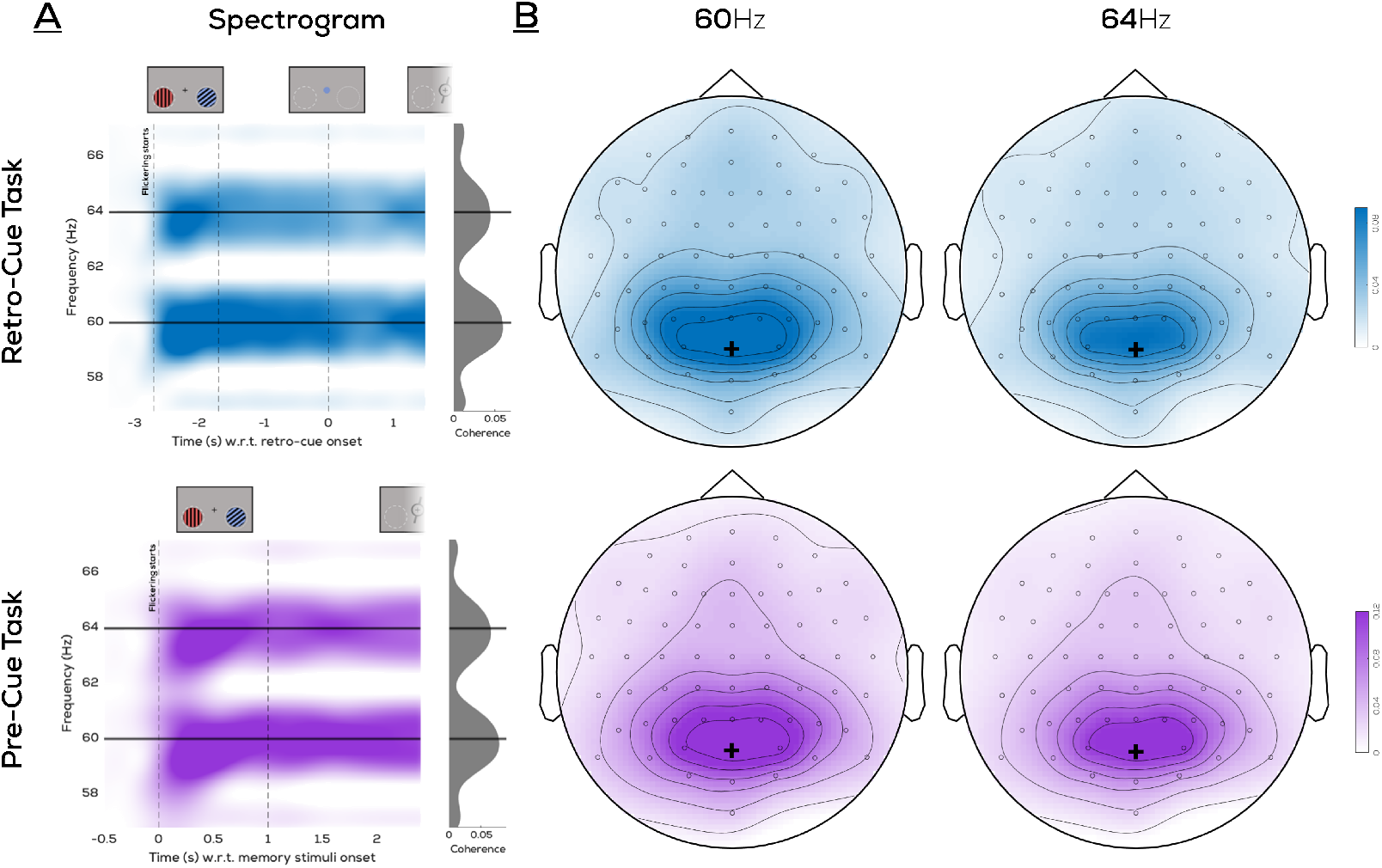
Average RIFT Coherence: **A:** Time-frequency plot averaged across participants and top 6 channels with highest coherence (selected individually per participant) showing clear peaks at 60Hz and 64Hz following flicker onset. **B:** Topographical distribution of average RIFT coherence over the 1s interval during which both gratings were on screen. Black plus marks the POz electrode.

We compared the RIFT responses (60 & 64 Hz averaged) elicited from locations of cued and uncued stimuli (Figure 3.2). In the retro-cue task, no difference between the RIFT signal evoked from the cued and uncued item locations was evident following presentation of the retro-cue. However, in the pre-cue task, a clear difference in coherence was observed between cued and uncued locations. A cluster permutation test (Maris & Oosterveld, 2007; green line in Figure 3.2) showed that in the pre-cue task, coherence from cued item locations was higher than that of uncued item locations in the interval from 0.28s before to 1.14s after stimulus onset (*p* < 0.0005; cluster t-mass compared to permutation distribution). The same test indicated no significant difference between cued and uncued item locations in the retro-cue task (*p* = 0.21; largest cluster t-mass compared to permutation distribution). We also compared the attentional modulation between both tasks by averaging coherence within the significant interval of the pre-cue task. This was done using a permutation test of mean differences (Ho et al., 2019), which confirmed that the attentional modulation (cued uncued coherence) in the pre-cue task was significantly stronger than that of the retro-cue task (mean difference = 0.023, 95% bootstrapped CI of mean differences = [0.013, 0.032], *p* < 0.0002). Though we present an average over coherence at 60Hz and 64Hz here, we also confirmed that this pattern of findings replicates for the 60Hz and 64Hz frequencies separately (Supplementary Figure S.1). In summary, we find enhanced processing at cued item locations during external shifts of attention but not during internal shifts of attention.

**Fig 3.2:**
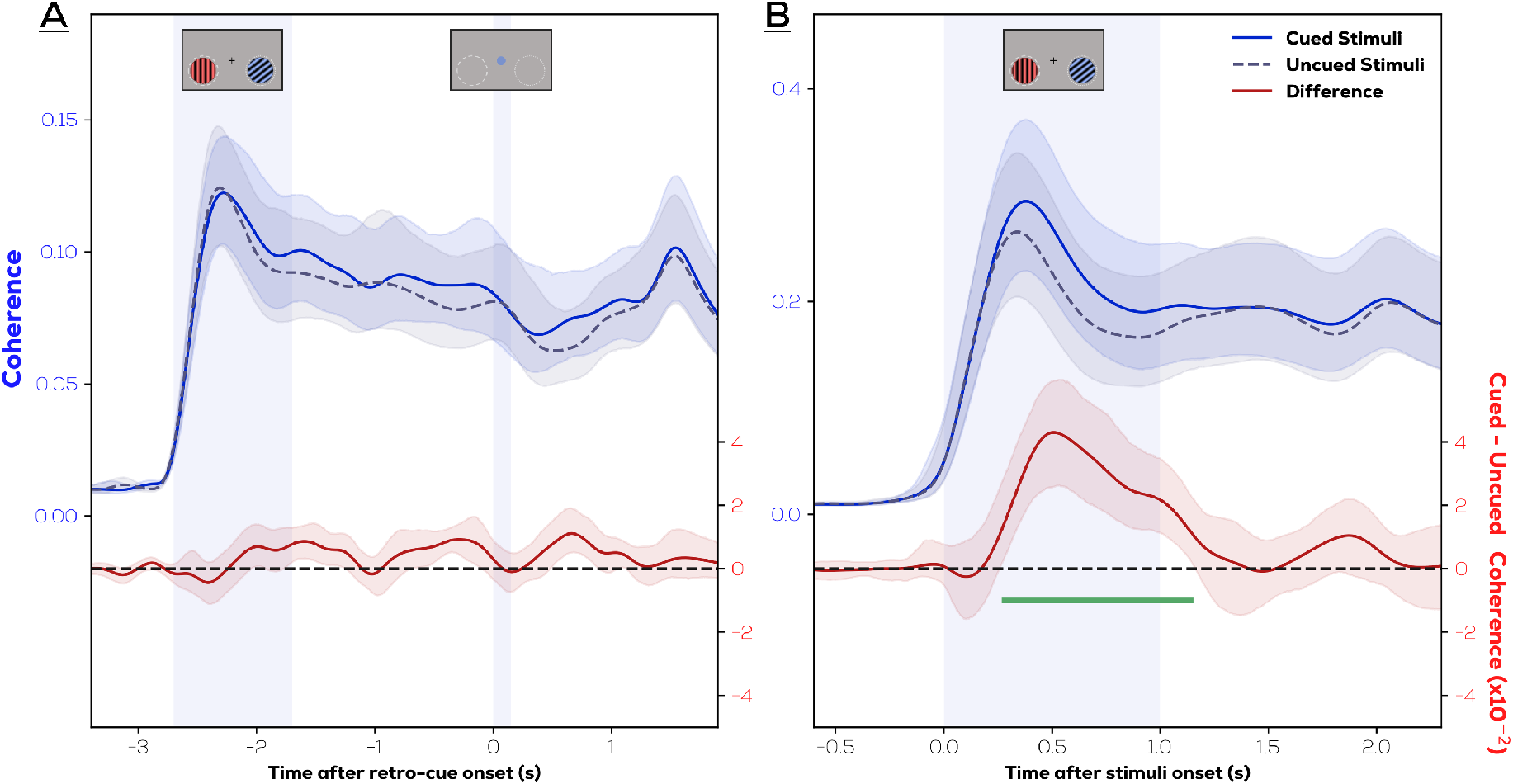
RIFT responses to prioritized locations are enhanced only during external and not internal shifts of attention. RIFT coherence from frequencies corresponding to cued vs. uncued stimuli (top) and difference (bottom) in the **A:** retro-cue task **B:** pre-cue task (shaded region - 95% bootstrapped CIs). Note that separate y-axis limits are used for the coherence traces over both figures to convey equivalent variances visually.

We next investigated potential lateralizations of alpha oscillations (8-13.5Hz) in both tasks. Desynchronization of alpha power contralateral to the previous/current location of the cued item has been identified as a marker of spatial attention (Ikkai et al., 2016; Liu et al., 2022; Poch et al., 2017; Sauseng et al., 2009). Both the retro-cue and pre-cue tasks showed posterior decreases in alpha power upon presentation of the retrocue or stimuli respectively compared to the baseline (Supplementary Figure S.2). Alpha lateralization is commonly associated with an attentional bias towards the cued item/location. The electrodes showing the strongest decrease in alpha power were contralateral to the direction of the cued item, confirming that both tasks showed strong alpha lateralization in parietal/occipital electrodes Figure 3.3A) (retro-cue task: mean = -1.021, 95% bootstrapped CI = [-2.02, -0.49]; pre-cue task: mean = -1.019, 95% bootstrapped CI = [-1.49, -0.68]). We compared alpha lateralization between the two tasks using a permutation test of mean differences (Ho et al., 2019), which revealed an equal degree of lateralization Figure 3.3B) across both tasks (mean difference = 0.193, 95% bootstrapped CIs of mean differences = [-0.14, 0.56]). This indicates that the previous location of the memorized item, although irrelevant for the task, was indexed by alpha oscillations, similar to what was observed during external attention. This confirms that people shifted attention (either to externally selected items or locations where internally selected items were previously memorized). Endogenous oscillations therefore suggest that internal and external selection operate with a similar spatial layout. This confirms that people shifted attention to the prioritized item in both our tasks and confirms earlier work suggesting that alpha oscillations index spatial attentional shifts (Ikkai et al., 2016; Kelly et al., 2006; Thut et al., 2006; Worden et al., 2000).

**Fig 3.3:**
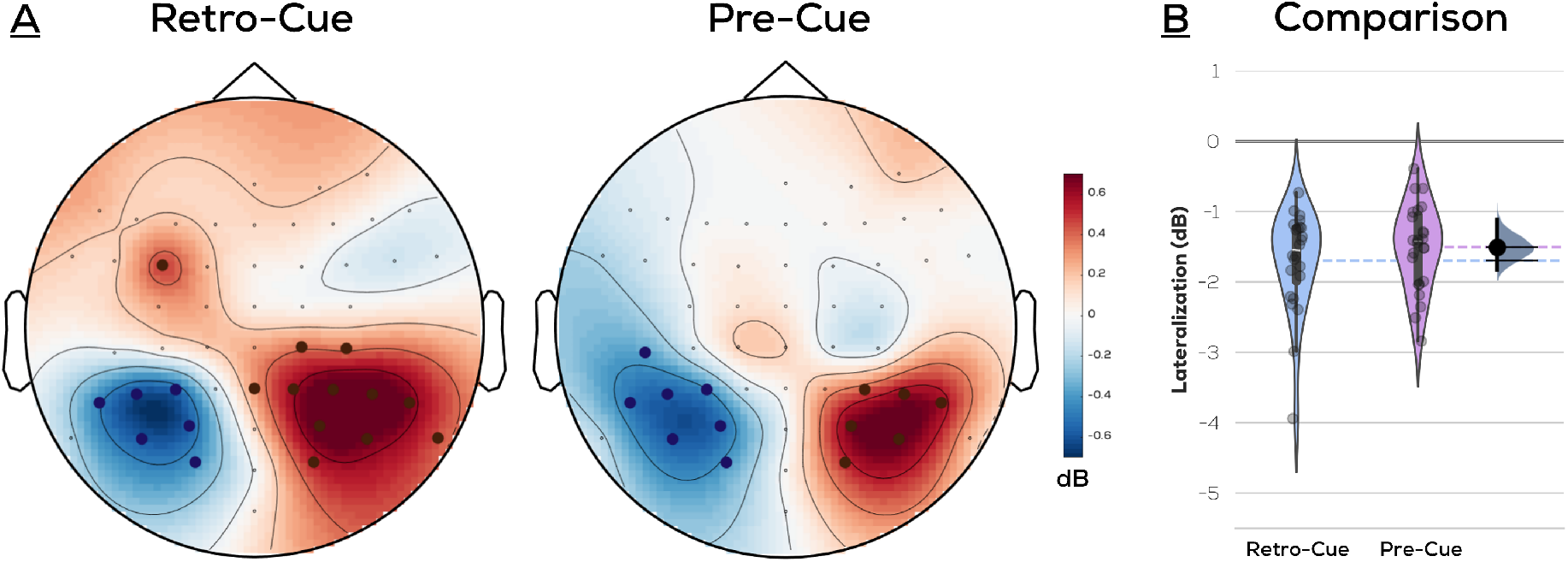
A: Alpha oscillations reflect the location (left/right) of the prioritized item during both internal and external selection. Average difference between alpha power (8-13.5Hz) in left and right item cued trials upon presentation of retro-cue/stimuli respectively (channel dots indicate 95% bootstrapped CIs excluding 0 and positive-red or negative-blue), **B: Comparison across tasks**. Shaded patch indicates kernel density estimation of respective scatterplots; dashed lines indicate respective means; gray indicates distribution of bootstrapped mean differences; black bar indicates 95% CIs.

We also looked at shifts in gaze position during both experiments, as these have been identified as correlates of prioritization in VWM. A cluster permutation test (Maris & Oosterveld, 2007; green line in Figure 3.2) was used to compare gaze position between left and right cued trials. Participants were instructed to maintain fixation throughout, and any trials where saccades were made to the item locations were excluded. Nonetheless, small deviations in gaze position were observed in the direction of the previous display location of the retro-cued item (*p* < 0.003; cluster t-mass compared to permutation distribution) in the retro-cue task (Figure 3.4A), and the cued item (*p* < 0.0005; cluster t-mass compared to permutation distribution) in the pre-cue task (Figure 3.4B). We also compared the gaze bias between both tasks by averaging coherence across time within the significant intervals. This was done using a permutation test of mean differences (Ho et al., 2019), which did not show any difference between the gaze biases between both tasks (mean difference = 0.043, 95% bootstrapped CI of mean differences =[-0.011, 0.125], *p* = 0.25). Thus, supplementing the results seen with alpha lateralization, small deviations in gaze position indexed the location where the cued item was previously memorized (retro-cue task) or shown (pre-cue task), despite this location being irrelevant for the retro-cue task.

**Fig 3.4:**
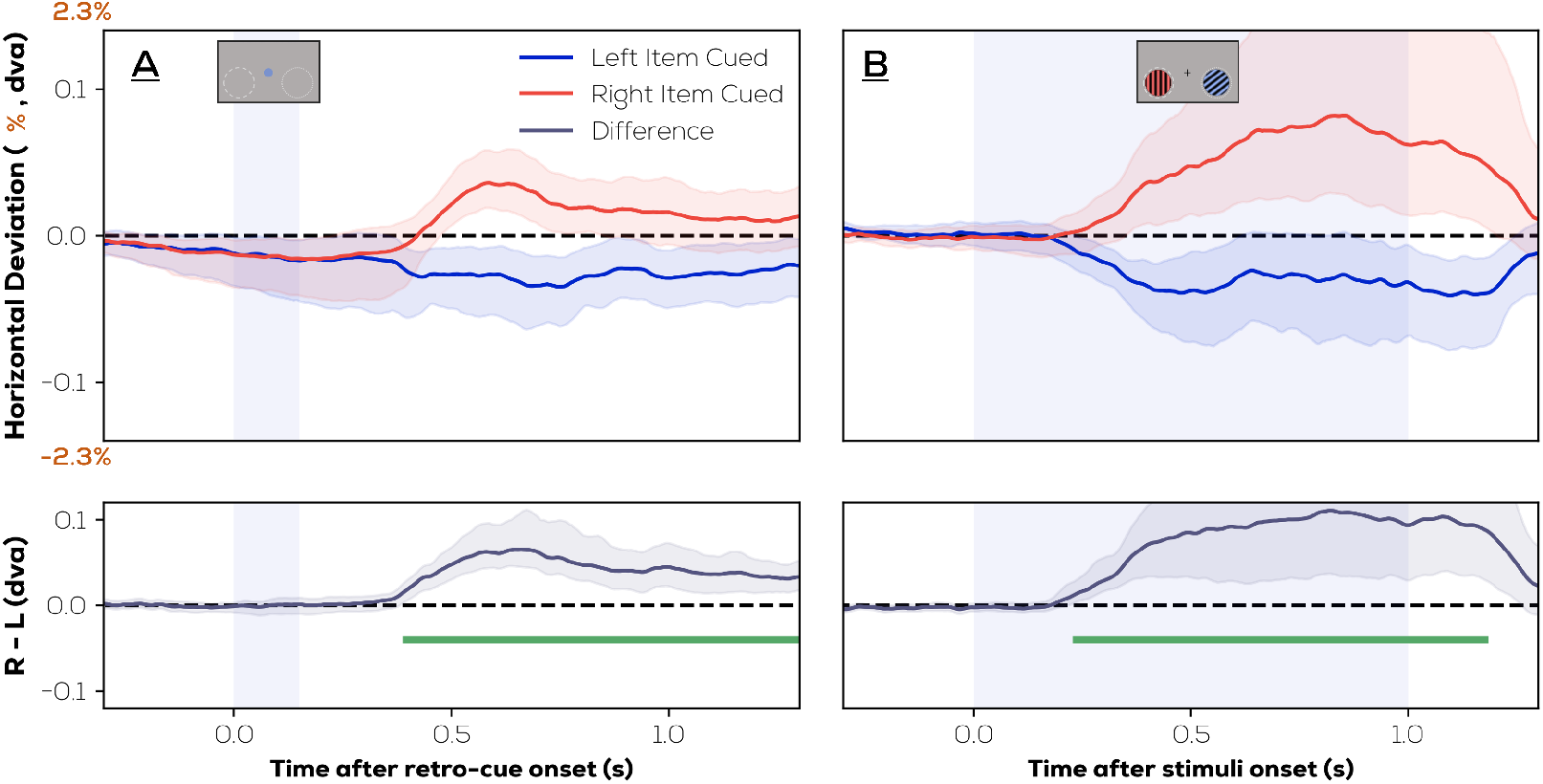
Mean horizontal gaze position across right cued and left cued trials (top) and difference (bottom) in the **A:** retro-cue task **B:** pre-cue task (shaded region - 95% bootstrapped CIs). Deviation is reported in terms of dva and % of distance from fixation to horizontal eccentricity of stimulus center.

Collectively, the alpha lateralization and the gaze bias results show substantial evidence of an attentional bias towards the location of the cued item, even in the retro-cue task when the memory items were no longer present.

The above results showcase three metrics (RIFT, lateralization of alpha power, and gaze bias) of attention. We ran linear mixed-effects models to investigate to what extent these metrics co-vary, and reflect similar or distinct cognitive mechanisms. We show that in both the retro-cue task where the RIFT response was not modulated by attention, as well as in the pre-cue task where it was, the trial-wise alpha lateralization and gaze biases did not predict the trial-wise RIFT response (Figure 3.5; top). This was the case despite accounting for variability across participants and the two frequencies used. The RIFT response is, however, inversely related to frequency (which reflects the fact that 64Hz tags evoke a weaker response than 60Hz tags), and in the pre-cue task is significantly higher from cued than uncued item locations (which reflects the result shown earlier Figure 3.2B with coherence). A second linear mixed-effects model showed that alpha lateralization is also not explained by either of the other two metrics in both tasks (Figure 3.5; bottom). This suggests that the three metrics reported here (namely RIFT, lateralization of alpha power, and gaze bias) capture distinct cognitive mechanisms.

**Fig 3.5:**
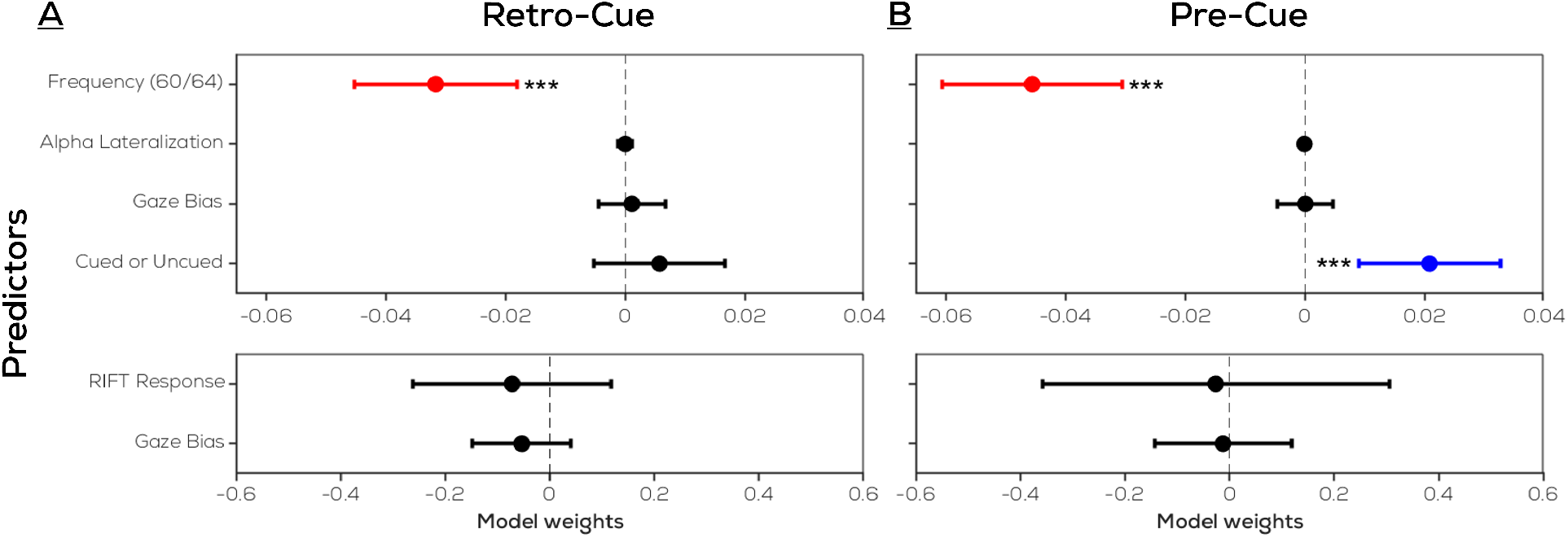
Gaze position bias does not drive other attentional modulations. Linear mixed effects model predicting top: the RIFT response, and bottom: alpha lateralization in the **A:** retro-cue task **B:** precue task (shaded region - 95% bootstrapped CIs). Linear coefficients (β) are reported with error bars representing 95% confidence intervals. The trial-wise gaze position bias does not explain the trial- wise attentional modulation in the RIFT response or alpha lateralization.

## Discussion

We investigated whether internal attention (orienting towards relevant content in VWM) boosts sensory processing at the attended item’s previous location, as is the case for external attention (orienting towards relevant stimuli in the environment). Despite a shift of attention towards the location of the prioritized items during both external and internal selection (as reflected by alpha oscillations and gaze position), early visual processing at locations of attended items is enhanced only during external attention. We thus show that the spatial organization within VWM is not a passive, vestigiallyderived consequence of internal and external attention utilizing the same neural machinery – which would have biased early visual processing in the same manner across the two. Our results instead suggest that this spatial organization uniquely facilitates their separate functions, and that the visual system uses space as an organizational principle to store and select items in VWM.

Our results are in line with other work providing evidence for a spatial organization of VWM. An established neural signature of attentional prioritization within VWM is the decreased amplitude of alpha oscillations observed in the hemisphere contralateral to the cued item, which we reproduced in both experiments (Liu et al., 2022; Poch et al., 2017; Sauseng et al., 2009). This lateralization indicates a spatial layout within VWM since it reflects the location where a prioritized item was previously memorized in the brain even when location is not required for the task. Furthermore, we show that participants’ gaze is biased toward the (now empty) location of the selected item, replicating existing work (de Vries & van Ede, 2024; Van Ede et al., 2019). The above results confirm that internal attention utilizes a spatial reference frame similar to that seen in external attention. Importantly however, this shared spatial bias does not extend to early sensory processing at the corresponding locations; although we observe a robust boost in sensory processing at cued item locations during external attentional selection, we do not observe this during prioritization in VWM. That is, though a substantial RIFT signal is evoked from the previous locations of prioritized and unprioritized items, these signals are not modulated by the retro-cue.

Behavioral work has shown that responses to targets are faster at previous locations of memorized items than other locations (Theeuwes et al., 2011). This is seemingly in contrast with our results, since we show that processing at these locations is not enhanced. Previous work (Theeuwes et al., 2011) has made use of perceptible stimuli presented at the relevant locations; participants responded to visible attentional probes faster at these locations than at others. We do not place any visible stimuli at these locations – this may be the differentiating factor. Responding faster to a target at a particular location does not guarantee that visual processing was already enhanced at that location before the stimulus was there. Here, we show that in the case of internal attention, the attended location in space does not evoke enhanced sensory processing. This follows logically from the fact that there is no task-relevant sensory information to process at the attended location. However, once a stimulus is shown in the corresponding location during the period of prioritization (Theeuwes et al., 2011), its processing could then benefit from the attentional bias that already exists more downstream along the visual hierarchy as captured by alpha lateralization (Liu et al., 2022; Poch et al., 2017; Sauseng et al., 2009). RIFT predominantly engages low level visual areas which would, in this case, not be engaged during VWM prioritization. There is evidence for this account: a bias in processing of the attended locations can be identified in a similar study using stimuli with visible, low-frequency flickers (Chota et al., 2024), and the previous location of the prioritized item can be decoded in early visual cortex when a visible outline is presented there (Zhou et al., 2022). Additionally, during spatial working memory, perceptual input traverses the visual system (in monkeys) faster when originating at the memorized location as compared to others (Roshanaei et al., 2024).

Overlap in areas utilized by external and internal processing is discussed within the sensory recruitment hypothesis. It posits that VWM storage and perception rely on equivalent patterns of neural activity in the visual cortex (Gayet et al., 2018; Ikkai & Curtis, 2011; Jerde et al., 2012; Scimeca et al., 2018). There is also evidence that features of prioritized VWM items are represented in sensory areas more strongly than those of unprioritized items (Christophel et al., 2018; Munneke et al., 2012), suggesting that neural populations representing prioritized VWM items and externally prioritized items overlap (Wan et al., 2020). It may be inferred from such accounts that internal and external attention rely on the same mechanisms. How does this fit with the lack of a prioritization reflected here in the RIFT response? A spatially selective boost in processing during internal attention need not follow directly from the fact that features of internally and externally prioritized items share patterns of activity in sensory cortices. It is possible that the similarities between internal and external attention at the neural level are present because prioritization in VWM accesses stored representations in sensory areas from a top-down direction. In the absence of any novel, perceivable sensory input to the system, this bias does not extend towards sensory processing at physical locations. Thus, our results are not in conflict with existing accounts of similarities between external and internal attention in sensory cortices; they simply show that although the neural mechanisms for representing visual contents during VWM and perception may be shared (i.e., sensory recruitment), the mechanisms for executive processes during VWM and perception (e.g., attentional selection) are not.

In tasks such as ours, the location of encoding is not relevant throughout the task. What use is there then in having this spatial organization in VWM at all? The visual system is constantly processing the world in a spatial manner; it is fine-tuned to parse items across some coordinate or location system. Therefore, whether necessary or not, it may be easiest to organize, store and retrieve VWM content with respect to this spatial coordinate system. This is akin to the “method of loci”, where items are supposedly better-remembered when mentally assigned to unique locations, or spatial contexts, which results in better memorization both in memory athletes as well as in naive participants upon training (Dresler et al., 2017; Wagner et al., 2021). There is support for such a spatial layout from evidence that locations (as compared to other visual features) play a special role in maintaining VWM; encoding VWM items at the same location instead of at unique locations decreases performance (Pertzov & Husain, 2014, but see Schneegans et al., 2021), feature bindings are stronger when locations are not shuffled (Treisman & Zhang, 2006), VWM performance is better for stimuli memorized on different depth planes rather than on the same depth plane (Chunharas et al., 2019), and the availability of location information before report improves memory performance more than the availability of orientation (Koevoet et al, 2024a, preprint) or color (Rajsic & Wilson, 2014) information. Thus, location has a special role among the features of content in VWM, and a spatial coordinate system is employed to better organize and recall memorized items.

Here, we use the RIFT response as a conduit for early sensory processing. Given the novelty of the key method used in this study, it is worth reflecting on this connection and on what the RIFT response actually represents. Using RIFT, we observe the response to rapidly flickering visual stimuli, the luminance of which changes too fast to be perceived but not too fast to evoke a neural response. It has been shown using source localization in MEG studies that the RIFT response predominantly reflects the earliest stages of visual processing at the level of V1 (Duecker et al., 2021; Ferrante et al., 2023), and that the topographic spread of brain areas responding to RIFT is a smaller and earlier subset of that forming the source of, for example, alpha oscillations (Ferrante et al., 2023). With this evidence on the RIFT response being localized specifically to early visual processing, our results are therefore not in conflict with previous work suggesting commonalities between external and internal attention more upstream in the visual hierarchy, such as the retrocue/stimuli evoked lateralization of alpha oscillations that we also observe here.

Since this is among the first studies to combine RIFT with EEG (Arora et al., 2024, preprint; Husta et al., 2024, preprint), the question may arise of whether we simply lack sensitivity to pick up attentional effects. We find this unlikely due to two reasons. First we show that attention does modulate the RIFT signal when directed at a perceived stimulus in our pre-cue experiment with identical stimulus parameters. The retro-cue and precue task maintained identical parameters in terms of the eccentricity and size of the flickering region. Second, even if no stimulus is presented at the tagged location, invisible background-flickers have been shown to evoke measurable neural responses that are modulated by attention (Brickwedde et al., 2022). These factors indicate that the method implemented here is sensitive enough to pick up on covert shifts of attention.

We also eliminated the possibility that the attentional RIFT modulation was actually caused by eye movements (which could bring the attended stimulus closer to the fovea, thereby enhancing the RIFT response), using linear mixed-effects analysis. Additionally, if RIFT modulations were explained by gaze position alone, we would also see an attentional modulation in RIFT for the retro-cue task where a gaze position bias is indeed visible. Notably, eye movements also did not predict alpha lateralization as shown previously (Liu et al., 2022).

Finally, we also show that the two neural markers of attention observed during shifts of external attention – namely alpha lateralization and the attentional RIFT modulation – do not co-vary. This is in agreement with previous work on alpha oscillations and frequency tagging (Gundlach et al., 2020; Zhigalov & Jensen, 2020). It has been suggested that modulations in alpha oscillations reflect a gating of visual information to downstream regions of the visual system (Rajsic & Wilson, 2014). This explains our observations: if the role of alpha oscillations is not to modulate excitability at the early sensory level, but rather to control the flow of information to regions further downstream in the hierarchy, it is not surprising that alpha lateralizations are observed here in the retro-cue task despite the absence of RIFT modulations. In the context of VWM, alpha oscillations are known to be load-dependent (Klimesch et al., 1999; Scheeringa et al., 2009; Tuladhar et al., 2007), reflecting an inhibition of visual input to better preserve memory content (Jensen & Mazaheri, 2010; Klimesch et al., 2007). Alpha lateralizations seen during internal attention may imply that preserving prioritized items in VWM requires alpha oscillations to control the spatial distribution of processing in downstream visual regions, without needing to spatially modulate the low-level sensory response. Thus, during prioritization in VWM, location is reflected at a level that includes alpha oscillatory activity, perhaps involving our internal representations of memorized objects, but not at a level that includes early visual processing.

## Conclusion

Here we investigated the role of early visual processing during attentional selection from perception and from visual working memory. We show that while both types of attentional selection operate within a spatial layout that reflects the spatial organization of our sensory environment, only external attention engages early visual processing. If a sensory boost was observed during internal attention, then its spatial layout may have been explained by internal attention vestigially displaying features of a mechanism that evolved for attending in the external world. By revealing a distinction between the mechanisms underlying internal and external attentional selection, our results show that internal attention is instead using location as an organizational principle to store and select items in VWM. We also show that although the neural mechanisms for representing visual contents during VWM and perception may be shared (i.e., sensory recruitment), the mechanisms for executive processes during VWM and perception (e.g., attentional selection) are not.

## Acknowledgements

This project receives funding from the European Research Council (ERC) under the European Union’s Horizon 2020 research and innovation programme (grant agreement n° 863732). We would like to thank Freek van Ede for valuable assistance in analyzing our eye movement data. We would also like to thank Laura van Zantwijk for assistance with data collection.

## Data and Code Availability

Raw and processed EEG/eye-tracking data, along with code corresponding to analysis and figures, are publicly available at https://osf.io/yrh6v/.

## Competing Interests

The authors declare no competing interests.

---

**Fig S.1:**
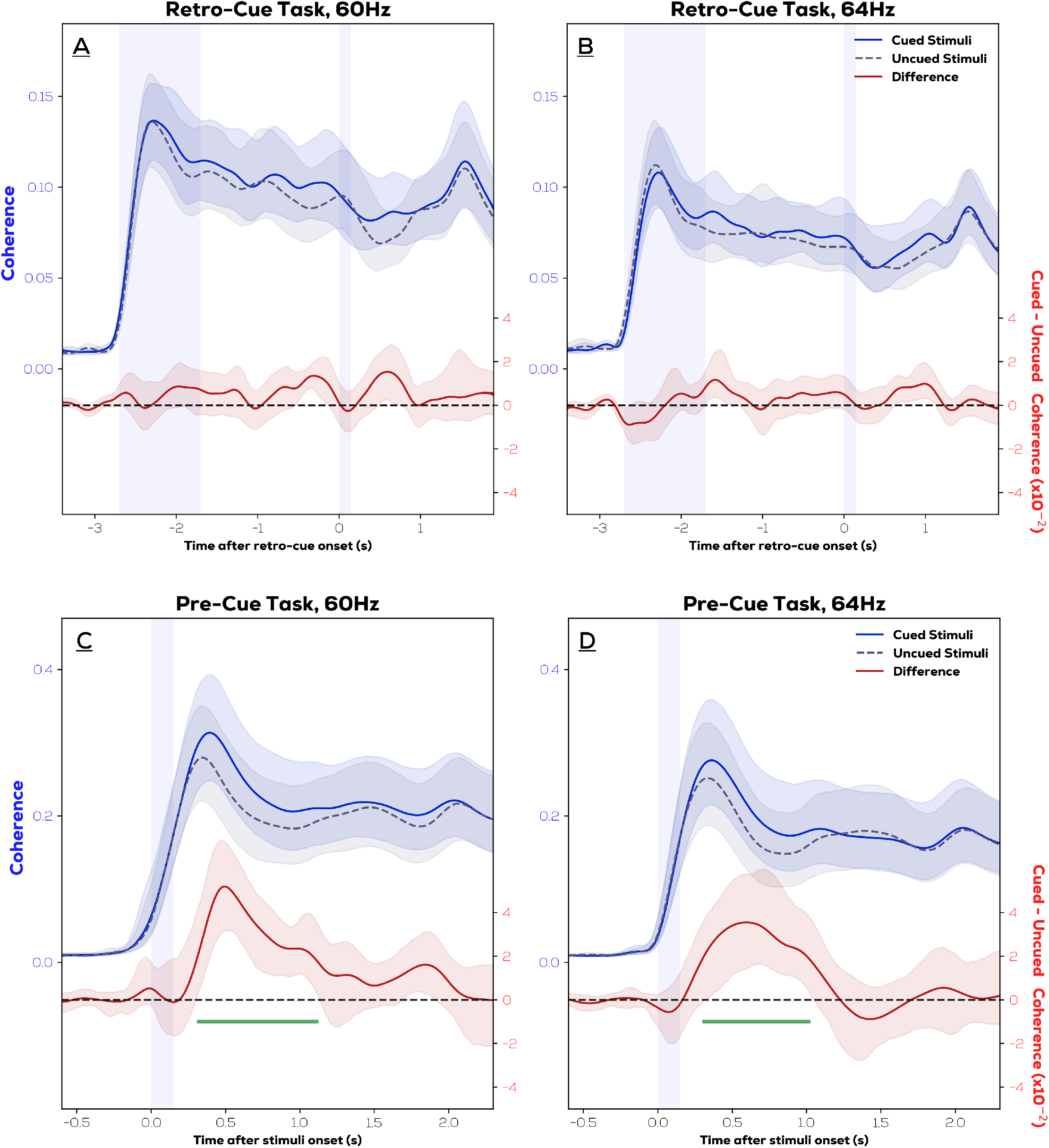
RIFT Coherence from frequencies corresponding to cued vs. uncued stimuli (top) and difference (bottom) in the **A-B**: retro-cue task **C-D**: pre-cue task (shaded region - 95% bootstrapped CIs). Extension to Figure 3.2, showing that the key effect reported in the main manuscript persists when separately analyzing responses to 60Hz and 64Hz tagged items.

**Fig S.2:**
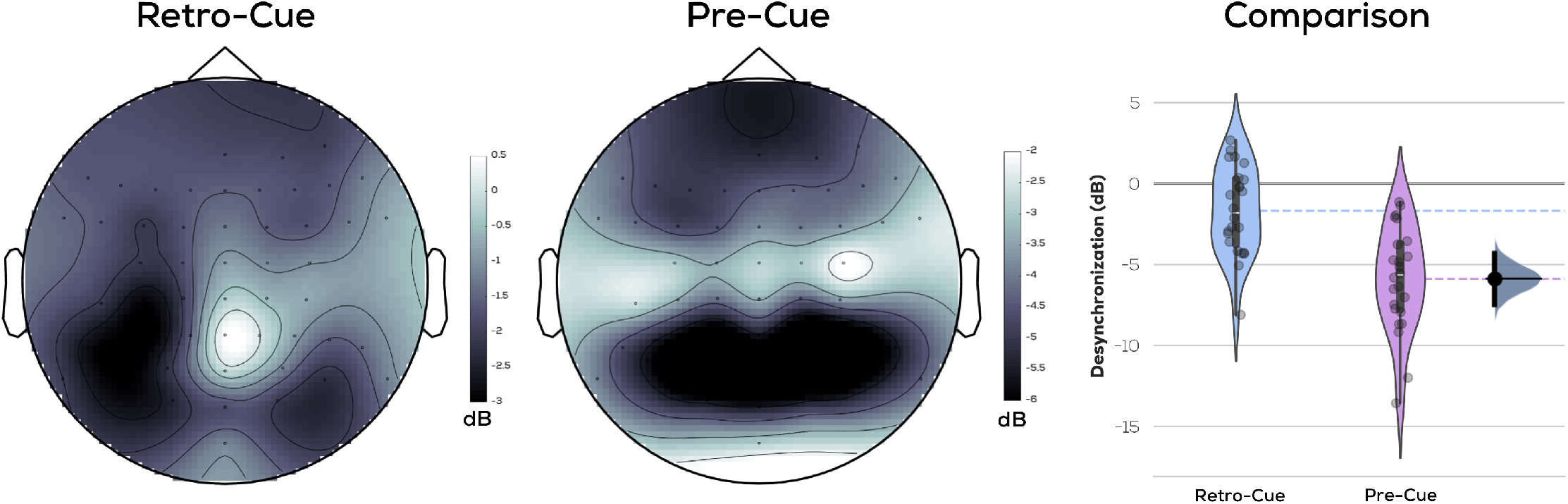
Alpha Desynchronization: In the manuscript we report lateralized alpha as a neural marker of spatial layouts in internal and external attention. Here, we show the alpha desynchronization – the average drop in alpha power (8-13.5Hz) upon presentation of retro-cue/stimuli – that was subtracted between left- and right-cued trials to produce the alpha lateralization, and its comparison across tasks using a permutation test of mean differences (Ho et al., 2019). (shaded patch indicates kernel density estimation of respective scatterplots; dashed lines indicate respective means; gray indicates distribution of bootstrapped mean differences; black bar indicates 95% CIs). The pre-cue task showed a considerably stronger decrease in alpha power (mean difference = -4.21, 95% bootstrapped CIs of mean differences = [-5.88, -2.61]).

## Notes

https://osf.io/yrh6v/.

### Competing Interest Statement

The authors have declared no competing interest.

